# An assessment of LRRK2 serine 935 phosphorylation in human peripheral blood mononuclear cells in idiopathic Parkinson’s disease and G2019S *LRRK2* cohorts

**DOI:** 10.1101/749226

**Authors:** Shalini Padmanabhan, Thomas A. Lanz, Donal Gorman, Michele Wolfe, Najah Levers, Neal Joshi, Christopher Liong, Sushma Narayan, Roy N. Alcalay, Samantha J. Hutten, Marco A. S. Baptista, Kalpana Merchant

## Abstract

The phosphorylated form of LRRK2, pS935 LRRK2, has been proposed as a target modulation biomarker for LRRK2 inhibitors. To qualify the biomarker for therapeutic trials, we assessed pS935 LRRK2 levels in Peripheral Blood Mononuclear Cells (PBMCs). Analyses of PBMCs from healthy controls, idiopathic Parkinson’s disease (iPD), and G2019S carriers with and without PD showed significant reductions in pS935 LRRK2 levels normalized to total LRRK2 levels in G2019S carriers with PD compared to those without PD or iPD. Neither analyte correlated with age, gender, or disease severity. Thus, pS935 LRRK2 in PBMCs may reflect a state marker for G2019S *LRRK2*-driven PD.

## INTRODUCTION

Missense mutations in the gene encoding leucine-rich repeat kinase 2 (*LRRK2*) have been associated with autosomal dominant Parkinson’s disease (PD) [1, 2]. The G2019S mutation in the activation loop of LRRK2 is the most common genetic cause of PD [3] and increases LRRK2 kinase activity, which is obligatory for the toxic effects [4]. Thus, discovery and development of LRRK2 kinase inhibitors is an actively pursued therapeutic strategy.

Direct assessment of LRRK2 activity has relied upon autophosphorylation of, serine 1292 (S1292) [5] or phosphorylation of its substrates, Rab8 and Rab10 [6]. However, the low stoichiometry of S1292 phosphorylation [5, 7] and non-specificity of Rab phosphorylation by LRRK2 [6] have hampered their utility as pharmacodynamic biomarkers. An alternative robust, but indirect, marker of LRRK2 activity is phosphorylation of serine 935 (pS935). Although the kinase responsible for phosphorylation of S935 remains an area of conjecture [8], all known LRRK2 kinase inhibitors reduce pS935 in cellular and animal studies [9, 10] indicating its usefulness as a pharmacodynamic biomarker of LRRK2 kinase inhibitors. However, since complex biological mechanisms appear to regulate phosphorylation/de-phosphorylation of S935 [11, 12] it is critical to qualify its utility in clinical samples from intended patient populations.

To this end, the Michael J. Fox Foundation for Parkinson’s research (MJFF) established the “LRRK2 Detection in PBMC Consortium”, a precompetitive alliance among MJFF and select pharmaceutical industry scientists. As a prerequisite to join the alliance, each member developed analytically validated assays for pS935 LRRK2 and total LRRK2 on a clinically viable platform along with associated assay performance data such as dynamic range, accuracy, precision, test-retest reliability and peripheral blood mononuclear cell (PBMC) collection protocols. After a head-to-head comparison of assays representing three different platforms (Meso Scale Discovery, Quanterix SIMOA single molecule array and CisBio homogeneous time-resolved fluorescence) using PBMCs collected under a standardized protocol at Columbia University Irving Medical Center (CUIMC), the SIMOA assay platform was selected for qualification studies.

This report describes assessment of total LRRK2 and pS935 LRRK2 in PBMCs collected from healthy controls, idiopathic PD (iPD) patients and G2019S *LRRK2* mutation carriers with and without PD.

## METHODS

### Subjects

Participants were recruited at CUIMC under an MJFF-funded LRRK2 biomarker project from March 2016 – April 2017. The study was approved by CUIMC institutional review board, and all participants signed informed consents. Participants underwent clinical evaluation using the Unified Parkinson’s Disease Rating Scale (UPDRS) and Montreal Cognitive Assessment (MoCA). Genotyping for G2019S *LRRK2* was conducted as previously described [13]. The demographics and clinical summary data on 117 study participants are presented in Table 1.

**Table 1.** Summary of demographic and clinical characteristics of the PBMC donors. The study cohort consisted of Controls (PD−) and PD subjects (PD+) with (+) and without (−) G2019S *LRRK2* mutation. SD = standard deviation; IQR = inter-quartile range; UPDRS = Unified Parkinson’s Disease Rating Scale.

| Characteristic | PD-G2019S-<br>(n=22) | PD-G2019S+<br>(n=16) | PD+G2019S-<br>(n=46) | PD+G2019S+<br>(n=33) |
| --- | --- | --- | --- | --- |
| **Age (years)** |  |  |  |  |
| Mean (SD) | 69.0 (7.1) | 57.9 (11.6) | 64.9 (9.4) | 71.7 (8.9) |
| Range | 57-85 | 37-83 | 41-82 | 56-91 |
| **Sex** | 13 M, 9 F | 6 M, 10 F | 29 M, 17 F | 19 M, 14 F |
| **Disease Duration (Years)** |  |  |  |  |
| Median (IQR) |  |  | 7 (2.25-10) | 11 (7-15) |
| Range |  |  | 0-25 | 0-26 |
| **Total UPDRS\*** |  |  |  |  |
| Mean (SD) |  |  | 28.1 (16.4) | 34.4 (18.7) |
| Range |  |  | 8-94 | 4-81 |
| **UPDRS Part 3\*** |  |  |  |  |
| Mean (SD) |  |  | 19.6 (12.3) | 22.2 (11.8) |
| Range |  |  | 5-64 | 1-53 |
| **MoCA\*\*** |  |  |  |  |
| Mean (SD) | 27.4 (2.0) | 28.9 (1.1) | 26.6 (3.0) | 26.1 (3.3) |
| Range | 23-30 | 27-30 | 12-30 | 18-30 |

\*UPDRS data available for only 64 PD subjects (81%). Total UPDRS includes UPDRS I-III.
\*\*MoCA data available for only 58 PD subjects (73%), and missing from 1 PD-G2019S-.

### PBMC Collection

Whole blood collected in PBS-diluted heparin was transferred into a Ficoll-filled Leucosep tube, centrifuged, upper phase extracted and centrifuged again. The supernatant was then decanted and cells washed. Cell counts were measured by a Countess Automated Cell Counter prior to resuspension and final centrifugation. The cell pellet was stored at −80°C until shipment.

### LRRK2 assays

PBMCs were lysed in the following buffer: 50 mM Tris-HCL, 150 mM NaCl, 1% Triton x-100, 2% Glycerol, 10 mM PPA, 20 mM NaF, 2 mM Na3VO4, 2 mM EGTA, and 2 mM EDTA at pH 7.5, with HALT protease and phosphatase inhibitors added fresh. 1 mL of cold lysis buffer was added to frozen aliquots of 3 million cells, and samples were sonicated for 5s on ice. Lysates were centrifuged at 14,000 x g for 3 min, supernatant was collected and stored at −80°C until day of assay. Samples were randomized on assay plates and blinded to group identity until statistical analysis. Total and pS935 LRRK2 assays developed on the Quanterix Simoa platform [14] used Neuromab N241A/34 for capture and biotinylated detection antibodies (Abcam clone UDD2 for total, and Cell Signaling clone D18E12 for pS935). Three technical replicates were run, and sample concentrations were interpolated from a standard curve ranging from 9.76 to 40,000 pg/mL of recombinant LRRK2 (Life Technologies A15197) in the following buffer (12 mM NaCl, 2.5 mM KCl, 1 mM MgCl_2_, 1.25 mM sodium phosphate, 2 mM CaCl_2_, 25 mM NaHCO_3_, 25 mM mannose at pH 8.13). Samples were diluted in the same buffer at 1:10 (total LRRK2) or 1:20 (pS935 LRRK2). The lower limit of detection for the assays was 19 pg/mL (total) and 4.2 pg/mL (pS935), and %CV for standard curves was 6%.

### Statistical Analysis

Pilot data collected on patient PBMCs informed the power calculations for the current study. Data were analyzed by Analysis of variance (ANOVA) with either two factors (all PD versus all controls) or four factors (PD+G2019S+, PD+G2019S−, PD−G2019S+, PD−G2019S−). For the latter, if the overall ANOVA showed p < 0.05, individual group differences were assessed by Student’s t-test. Sensitivity analysis adjusting for age and sex were conducted using linear regression. Correlation coefficients were based on Pearson correlation coefficients that included all data.

## RESULTS

Total and pS935 LRRK2 levels were first compared for all PD versus all control subjects regardless of G2019S *LRRK2* mutation status. While the variances appeared to be higher for the PD subjects (standard deviation = 4,863 vs. 3,084), there were no significant differences between PD and control groups for either total LRRK2 (p = 0.18) or pS935 LRRK2 levels (p = 0.92) whereas the ratio of pS935 to total LRRK2 showed a significant decrease in PD cases versus non PD controls (p=0.007). When controls and PD subjects were segregated by the G2019S mutation status, there were no difference in total LRRK2 levels among the 4 groups (p=0.13, Figure 1A). However, ANOVA indicated a significant group effect for pS935 LRRK2 (p=0.028), with PD+G2019S+ subjects showing ~32% lower pS935 LRRK2 levels than PD+G2019S- subjects (p<0.05, Figure 1B). Interestingly, the ratio of pS935 to total LRRK2 was significant across the groups by ANOVA (p < 0.01) with the PD+G2019S+ group showing significant reductions in the ratio when compared to all other groups (p<0.05; Figure 1C).

**Figure 1.**
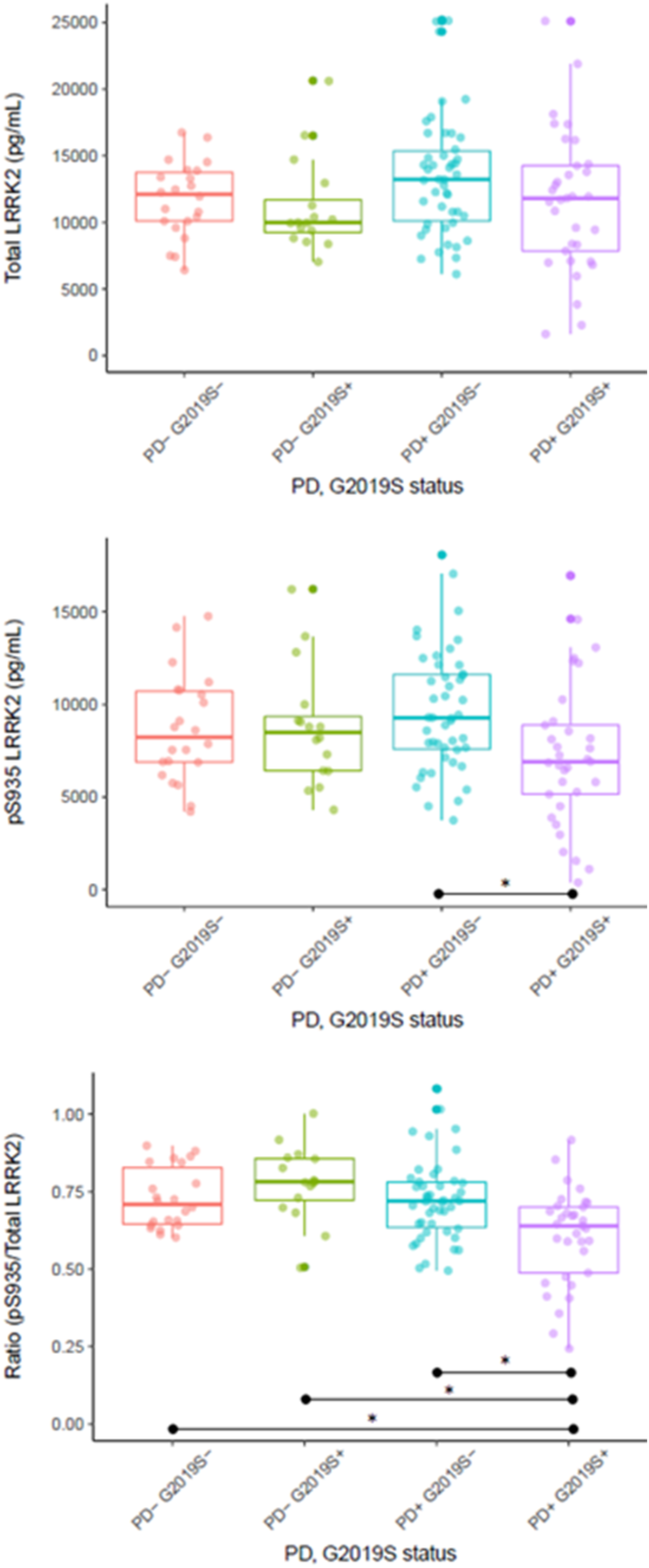
Total and pS935 levels in PBMCs. Total LRRK2 and pS935 levels are shown in panels (A) and (B), respectively whereas panel (C) depicts the ratio of pS935 to total LRRK2. * group differences at p<0.05.

The LRRK2 measures were compared to Montreal Cognitive Assessment (MoCA) data and UPDRS scores (available for 94% and 83% of subjects, respectively) as well as to the sex of participants, age at donation of the biospecimens and disease duration (where applicable). Although there was a strong correlation between total and pS935 LRRK2 levels (r = 0.89), total and pS935 LRRK2 levels did not correlate with any demographic or clinical factors (-−0.25 < r < 0.25, data not shown).

## DISCUSSION

In this study we demonstrate an unanticipated decrease in pS935 LRRK2, but not total LRRK2, levels in human PBMCs from PD subjects with the G2019S *LRRK2* mutation compared to iPD subjects, using a sensitive and robust digital immunoassay. Moreover, when normalized to total LRRK2, statistically significant decreases in pS935 LRRK2 were apparent not only between disease-manifesting G2019S *LRRK2* carriers and iPD subjects but also between the former group and non-manifesting G2019S *LRRK2* carriers as well as healthy controls. Since G2019S mutation results in higher LRRK2 kinase activity [4], the decrease in pS935 LRRK2 levels in carriers with PD was unexpected and indicates the presence of distinct biological mechanisms in LRRK2 G2019S mutation-driven PD compared to iPD or non-manifesting carriers. However, pS935 LRRK2 levels did not correlate with disease severity (based on UPDRS or MoCA), disease duration, age at biospecimen donation, or other demographic factors.

There is only one previous report [15] on levels of pS935 and total LRRK2 in PBMCs, but these assessments were restricted to iPD and healthy controls. Consistent with our data, they reported no differences in pS935 or total LRRK2 levels in iPD subjects compared to healthy controls. On the other hand, Atashrazm et al. [15] reported higher levels of total LRRK2 levels in neutrophils from iPD cases compared to healthy controls. The apparent discrepancy between our study and [15] may be due to previously reported cell-specific regulation of LRRK2 expression [16,17].

Although PBMCs represent a relatively simple biospecimen collection method, future studies of total and pS935 should examine distinct cell types such as monocytes and neutrophils since LRRK2 expression is dramatically higher in these cells [16] and the cellular heterogeneity of PBMCs may have masked alterations in total or pS935 LRRK2. Another major caveat of the present study is that we did not assess direct markers of LRRK2 kinase activity such as pS1292 LRRK2 or pT73 Rab10 [6]. In future studies, it will be critical to assess these markers along side pS935 LRRK2 levels to study whether despite higher kinase activity of G2019S LRRK2, compensatory biological mechanisms lead to de-phosphorylation of the S935 residue.

In summary, the data presented here indicate that normalized pS935 LRRK2 level has the potential to be a disease state biomarker of G2019S *LRRK2*-associated PD but the results need to be replicated and extended in an independent and larger cohort. The unexpected decrease in pS935 LRRK2 levels in G2019S-positive PD is indicative of distinct biological mechanism(s) at play in this patient population and underscores the importance of collaborative consortia for biomarker qualification studies using standardized human biospecimen collections.

## ACKNOWLEDGMENTS INCLUDING FUNDING SUPPORT

The Michael J. Fox Foundation would like to thank all the members of the Industry LRRK2 Detection Consortium (Omar Mabrouk from Biogen, Sarah-Huntwork Rodriguez from Denali Therapeutics, Matthew Fell, Julie Lee and Payal Sheth from Merck & Co., Inc., Thomas Lanz from Pfizer Inc., Nathalie Schussler from Sanofi and their respective teams) for their participation and engagement in the Consortium initiative.

SIMOA assay development and testing was funded by Pfizer Inc. Roy N. Alcalay received research funding from the NIH, Parkinson’s Foundation and the Michael J Fox Foundation.

